# Perceiving animacy in ‘identical’ images

**DOI:** 10.64898/2026.04.01.715738

**Authors:** Tal Boger, Chaz Firestone

## Abstract

Some objects appear ‘animate’ (e.g., dogs and elephants) while others do not (e.g., boots and sofas). This distinction pervades human cognition, with an expansive literature reporting striking effects of animacy on vision, memory, social perception, and neural organization. But studies of perceived animacy face a persistent challenge: Objects that differ in animacy tend to differ in many lower-level visual features (e.g., shape, texture, spatial frequency). Thus, it remains controversial whether animacy per se — as opposed to its lower-level correlates — drives visual processing. Here, we achieve previously unattainable levels of experimental control to demonstrate that the visual system represents animacy itself, beyond its lower-level covariates. We vary animacy while holding nearly all lower-level features constant by exploiting “visual anagrams” — a diffusion-based technique for generating static images whose interpretations change radically with orientation. Eight pre-registered experiments leverage this approach to demonstrate that representations of animacy structure visual working memory and guide visual attention. Thus, the visual system extracts animacy itself, beyond its lower-level correlates.

## Main text

Among the most consequential distinctions in human cognition concerns an object’s *animacy*: Even when motionless, some objects appear animate (e.g., tigers and bears), while others do not (e.g., rocks and sticks). Representing animacy is of critical importance, helping us recognize threats, find friends or caretakers, predict behavior, and more generally determine which entities in the world have beliefs, desires, and goals. Indeed, researchers across many disciplines have hypothesized that animacy is an ancestrally prioritized feature that may be privileged in the mind [1]. Accordingly, an expansive psychological literature has catalogued numerous effects of perceived animacy on various mental processes, including visual search [2], change detection [1], working memory [3], word learning [4], and even neural organization [5-7]. For example, it is easier to locate an animate object (e.g., a dog) among inanimate objects (e.g., tables, plates, and chairs) than among other animate objects (e.g., birds, fish, and cats), suggesting that basic processes of visual attention are sensitive to the animacy of visual stimuli [1,8].

However, all such research faces a challenge: Objects that differ in animacy also differ in a variety of additional features, including shape, curvature, texture, and other ‘low-level’ stimulus properties. Thus, it is difficult to attribute experimental effects to animacy itself as opposed to these lower-level covariates. Indeed, previous work suggests that systematic differences in curvature explain away effects of animacy on search [8], and that shape and texture alone may be sufficient to classify animates [9,10]; such regularities even apply to studies of perceived animacy in macaques [11]. Though some approaches show promise in disentangling animacy from one correlate at a time [12], a comprehensive and generalizable solution remains an unrealized goal.

### The present work: Perceived animacy in visual anagrams

Here, we pursue a new approach to studying perceived animacy. Recent advances in generative artificial intelligence enable the creation of ‘visual anagrams’ [13] — images whose interpretations change from one category to another when rotated. We generated a novel stimulus set of anagrams in ways that varied animacy — for example, a dog in one orientation and a boot when rotated, or an elephant in one orientation and a sofa when rotated. Whereas natural images (or line drawings) of dogs and boots differ in animacy, they also differ in shape, curvature, and texture. However, when a visual anagram looks like a dog in one orientation but a boot in another, it is *the very same image* in both cases — composed of the same pixels, just rotated (Figure 1) — such that nearly all lower-level features remain constant (see Materials and Methods for further discussion) while animacy changes categorically. In this way, visual anagrams enable far greater control of high-level perceptual properties than has previously been possible [14]. Eight pre-registered experiments leverage this approach to isolate perceived animacy, demonstrating effects on perception, attention, and working memory.

**Fig. 1.**
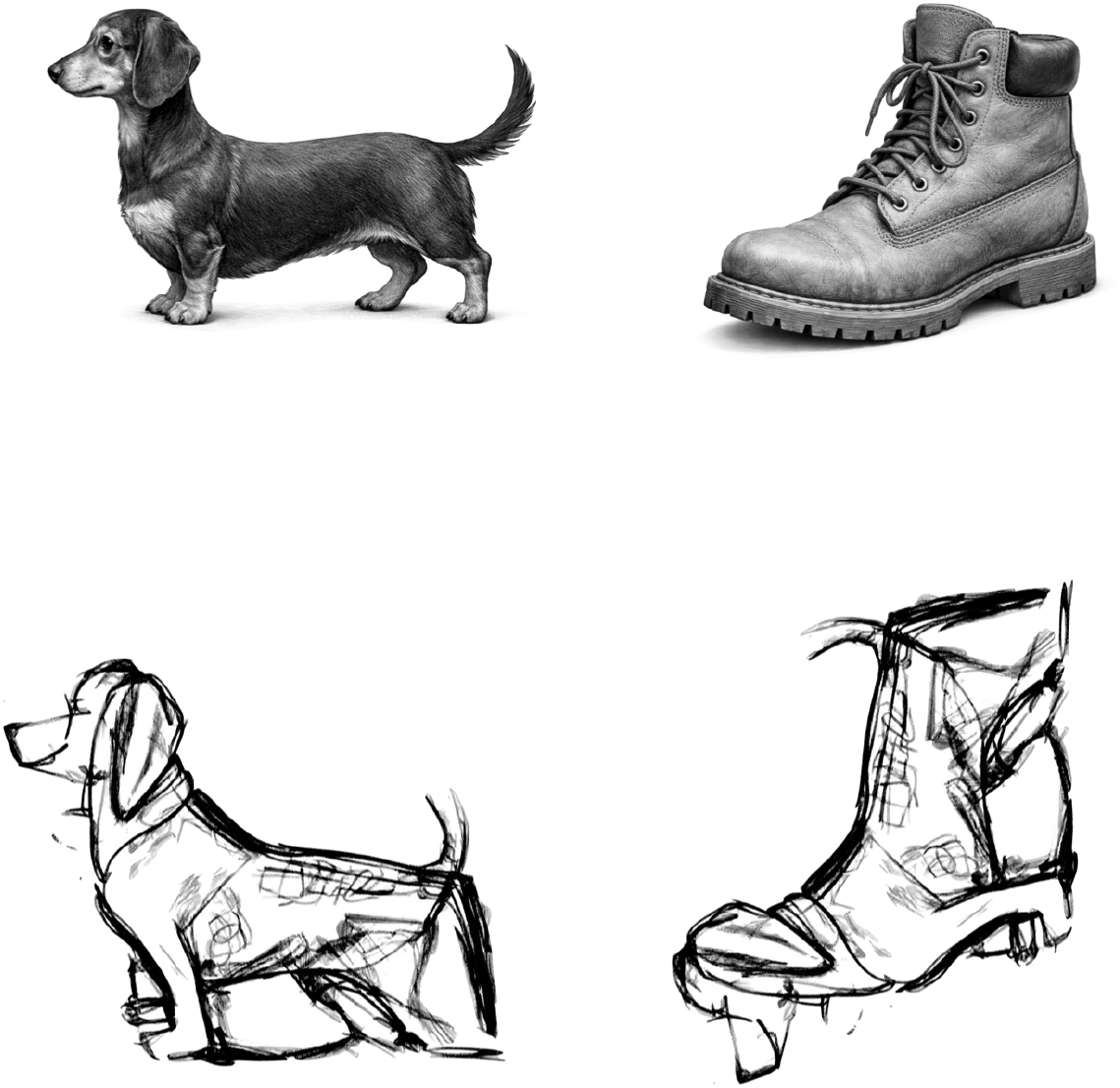
(Top Row) This dog and boot differ in animacy, but they also differ in many lower-level features. (Bottom Row) This dog and boot are visual anagrams: They differ in animacy while preserving nearly all lower-level features, because they are the very same image rotated 90°.

## Results

Experiment 1 used a visual working memory task (Figure 2A). On each trial, participants viewed an array of 5 unique anagrams (e.g., airplane, horse, rabbit, elephant, car). After a 3-second retention interval, one of the images was replaced with a new anagram that was held out of the original 5 (e.g., dog/boot), and participants had to indicate which item had changed by clicking on its position. Crucially, changes either altered the item’s animacy (e.g., rabbit→boot) or did not (e.g., rabbit→dog). We observed higher accuracy for changes that altered animacy vs. those that did not (*M* = 3.7%, *t*(27) = 2.36, *p* = 0.025, *d* = 0.45); in other words, rabbit→boot was more detectable than rabbit→dog, even though (a) the boot and dog are the very same image (just rotated), and (b) animacy itself was completely task-irrelevant (and was never mentioned to subjects).

**Fig. 2.**
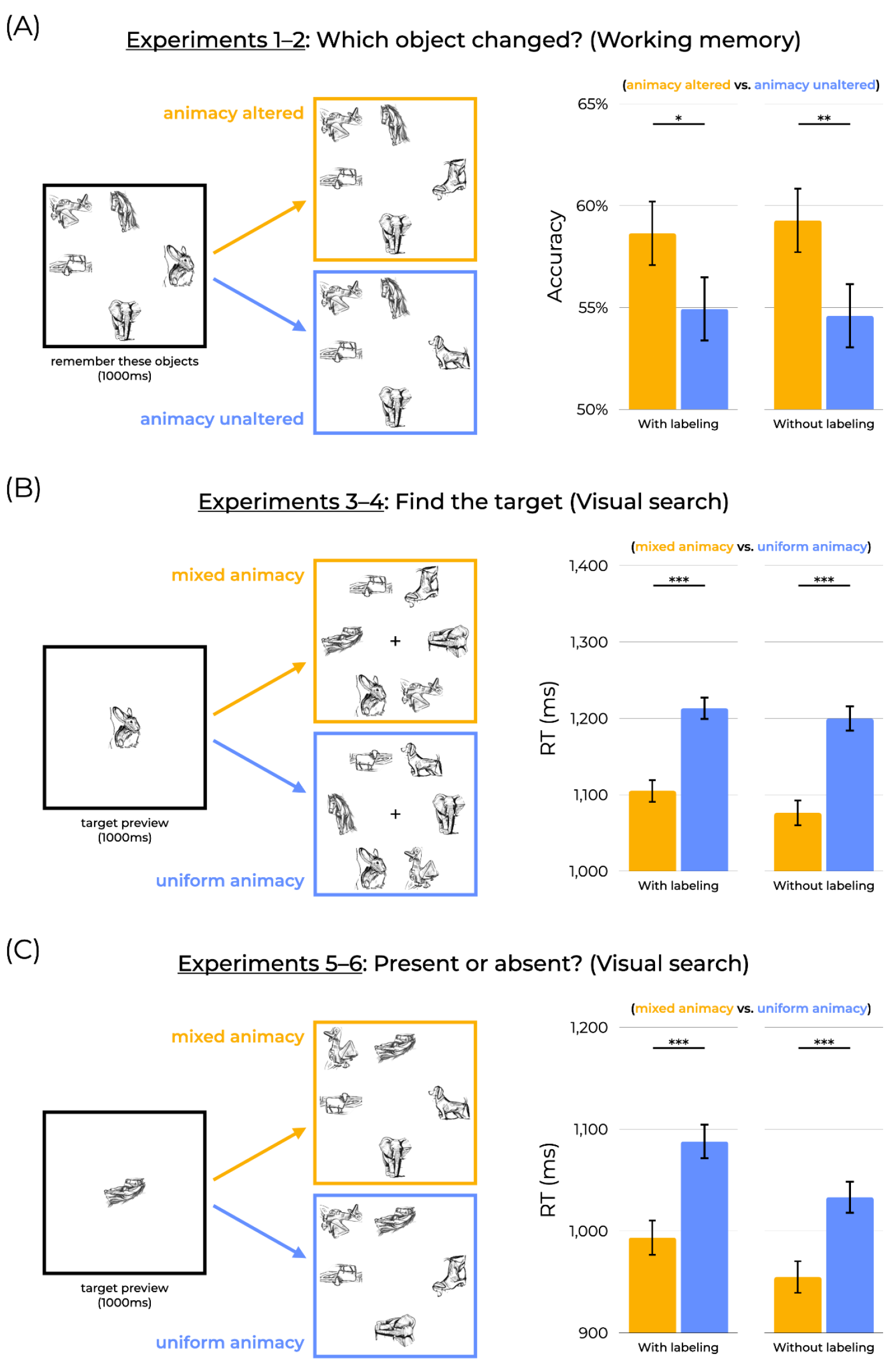
(A) In a visual working memory task, changes that altered animacy (e.g., rabbit→boot) were more detectable than changes that did not (e.g., rabbit→dog), even though animacy was task-irrelevant. (B) In a visual search task (modeled after [2]), participants were faster to find targets in mixed animacy arrays than in uniform animacy arrays. (C) This search effect also arose in a present/absent search task. Error bars represent standard errors of the differences across conditions. Readers may view our tasks at https://perceptionresearch.org/anagrams_animacy.

Importantly, Experiment 1 contained a ‘labeling’ phase where participants viewed all the anagrams and matched them to their corresponding labels before the start of the task (to ensure the anagrams were recognized as intended). Experiment 2 repeated Experiment 1 without this phase, and found the same pattern: Participants were more accurate for changes that altered animacy than those that did not (*M* = 4.7%, *t*(27) = 2.99, *p* < .01, *d* = 0.57), internally replicating our key result and ensuring that the labeling phase did not exert undue bias on participants.

Might animacy also affect more canonically *visual* processes, such as attention? This possibility is contested in the literature; whereas some studies find effects of animacy on visual search [2], others suggest that facilitated search is driven merely by differences in curvature between animates and inanimates [8]. In Experiments 3–4 (Figure 2B), we adapted a visual search task from Long et al. (2017). On each trial, participants previewed one anagram image (their target) and then saw an array of 6 unique anagrams; the task was simply to locate the previewed target within this array (and indicate its location by pressing a number key corresponding to its position). Half of trials contained ‘uniform animacy’ search arrays (where all objects were either animate or inanimate), while the other half contained ‘mixed animacy’ arrays (where an animate target was surrounded by inanimate distractors, and vice versa). We observed faster search on mixed animacy trials (*M* = 108ms, *t*(28) = 7.04, *p* < .001, *d* = 1.31; Experiment 3) — even though the mixed animacy and uniform animacy trials contained the exact same images (just rotated). As before, this effect replicated without the labeling phase (*M* = 124ms, *t*(29) = 7.10, *p* < .001, *d* = 1.30; Experiment 4).

Experiments 5–6 (Figure 2C) further tested effects of animacy on basic mechanisms of visual attention by using a more conventional visual search task. Here, participants indicated whether a target object was present or absent in a search array (uniform animacy or mixed animacy). As before, search was faster on mixed animacy trials than uniform animacy trials (*M* = 94.4ms, *t*(29) = 5.37, *p* < .001, *d* = 0.98; Experiment 5) — an effect that also held without the labeling phase (*M* = 78.5ms, *t*(28) = 4.80, *p* < .001, *d* = 0.89; Experiment 6).

In each of the above cases, our observed effects cannot be attributed to the lower-level features researchers typically worry about in similar studies (e.g., shape, texture, etc.), because orientation is the only feature that changed across the two interpretations of each anagram. But changes in orientation are of course changes to an image’s low-level features; what if that difference explains our results? In other words, perhaps our previously observed effects merely reflect search advantages for orientation, aspect ratio, or center of mass. Though there is little independent reason to suppose that animates and inanimates are, as a group, distinguished by orientation or aspect ratio, we nevertheless addressed this issue in a follow-up experiment. Experiment 7 (Figure 3A) repeated the present/absent search task from Experiments 5–6 but contained a block of trials that depicted silhouetted versions of the anagrams (via convex hull). These silhouettes are just indistinct blobs (and thus not identifiably animate or inanimate) but they share orientation and aspect ratio information with our original stimuli; thus, they reveal whether orientation alone produces similar effects. We found no search advantage for the silhouetted stimuli (*M* = 2.16ms, *t*(144) = 0.17, *p* = 0.86, *d* = 0.01, BF10 = 0.09). Crucially, Experiment 7 also included anagram trials of the sort appearing in Experiments 5–6. This allowed us to replicate (once again) the key animacy effects with visual anagrams (*M* = 75.7ms, *t*(144) = 6.55, *p* < .001, *d* = 0.54), and perhaps most importantly ensure that the *difference* in search advantage between the two stimulus classes (anagrams vs. silhouettes) was itself significant (*M* = 73.5ms, *t*(144) = 4.24, *p* < .001, *d* = 0.35). This last result ensures that even if there were a small but undetected role for orientation, our effects go beyond its contribution.

**Fig. 3.**
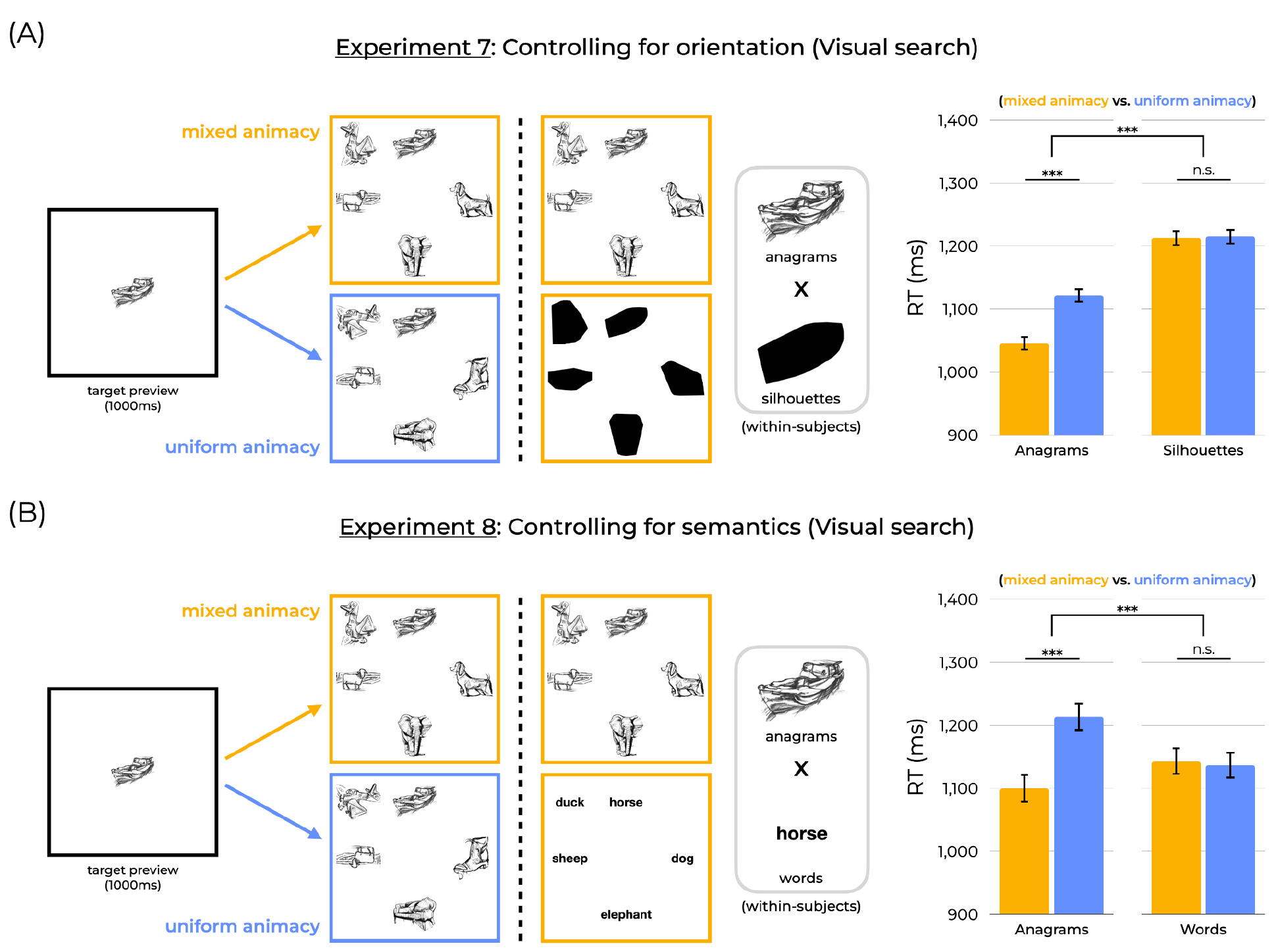
(A) To control for orientation (perhaps the only feature left uncontrolled by visual anagrams), we replicated the present/absent search task with silhouetted versions of the anagrams (within-subjects). While anagrams again showed a search advantage, silhouettes did not, suggesting that orientation differences cannot account for our results. (B) To control for linguistic labeling and/or effects of semantics, we replicated the present/absent search task once again with linguistic labels for the anagrams (within-subjects). We observed a significant search advantage for the anagrams, but not for the words, meaning that linguistic labeling cannot account for our results. Error bars represent standard errors of the differences across conditions.

In a final experiment (Experiment 8; Figure 3B), we asked whether our search effects merely reflect semantic labeling processes. In other words, linguistic labels of the depicted objects access their semantic representations of animacy; would simply replacing the anagram images with their linguistic labels produce the same search effects? Experiment 8 used the same design as in Experiment 7 but with written labels. Participants completed a present/absent search task. One half of trials presented anagram images, and the other half presented labels (i.e., the text “dog” and “boot”, etc., appearing in a search array). Once again, we replicated the previous search advantages for anagrams (*M* = 114ms, *t*(29) = 5.04, *p* < 0.001, *d* = 0.92). However, we observed no search advantage for the words (*M* = −6.38ms, *t*(29) = 0.30, *p* = 0.77, *d* = 0.05, BF10 = 0.20); and the difference in search advantages between the anagrams and words was itself significant (*M* = 120.0ms, *t*(29) = 3.68, *p* < 0.001, *d* = 0.67). Given that linguistic labels for the anagrams fail to produce a significant search effect, our results implicate visual processing of animacy rather than merely any semantically activated representation of animacy.

## Discussion

Is visual processing sensitive to animacy itself? Given the centrality of animacy to a range of cognitive processes, this question has motivated much research in psychology and neuroscience. However, it has remained without a conclusive answer because animacy is tightly (and seemingly intrinsically) entangled with shape, curvature, texture, and other lower-level features. The present work takes a novel approach to this question, finding that even when two images are pixelwise identical in ways that isolate animacy from its lower-level covariates, they are processed differently by basic processes of perception and cognition.

Our approach builds upon related work exploring the relation between lower-level features and high-level properties. For example, one popular approach involves ‘texforms’, a texture-synthesis algorithm that transforms stimuli into unrecognizable blobs, disrupting high-level recognition but preserving mid- and low-level features [2,6,15]. Our approach provides a complement, isolating animacy from its lower-level correlates rather than the other way around.

Our findings are also consistent with theoretical frameworks emphasizing the importance of animacy in human cognition. Such theories suggest that the centrality of animacy in our lives (both in modern times and ancestrally) has led the mind to prioritize it, producing effects of animacy on basic cognitive tasks including visual change detection [1,16]. Our work offers even stronger and more controlled evidence for many of these conclusions.

Although our approach rules out alternative, lower-level explanations, it remains possible that a high-level property other than animacy — such as natural vs. manmade, which also categorically distinguishes our stimuli — might explain our results. This possibility would still appeal to a (quite related) high-level property and so remain consonant with our general claim. Still, future work could nevertheless address this question directly; for example, rocks, trees, and lakes are natural but inanimate, and so could serve as stimuli in similar experiments.

Finally, the present work interfaces with both longstanding and contemporary philosophical discussions of high-level perception. As Block [17] writes: “What is a high-level representation? Is it just a matter of associations among low-level properties?”. Our work suggests that the answer to this latter question is “no”. At least for the case of animacy, high-level perception can be disentangled from its lower-level correlates.

## Methods

Supplementary methods appear in SI Appendix. Studies were approved by the Johns Hopkins Institutional Review Board; participants gave informed consent.

## Data, Materials, and Software Availability

Anonymized participant data, along with pre-registrations, experiment code, and analysis scripts are available on the Open Science Framework (https://osf.io/rw8tu/).

## Materials and methods

### General Methods

#### Open Science Practices

All experiments reported in this paper were pre-registered before data collection. Pre-registrations cover experimental design, sample size, exclusion criteria, and analysis plans. Interested readers may view all experiments at https://perceptionresearch.org/anagrams_animacy. An OSF repository containing all data, pre-registrations, analysis scripts, and experiment scripts is available at https://osf.io/rw8tu/.

#### Participants

All participants were recruited via the online platform Prolific (for a discussion of the reliability of this participant pool, see Peer et al., 2017). Each study recruited unique participants.

#### Stimuli

We used the “visual anagrams” model presented in Geng et al. (2024) to generate our stimuli. (Images were generated via cluster jobs on the Advanced Research Computing at Hopkins core facility.) Stimulus generation proceeded as follows: First, we created a list of animate and inanimate object labels. Then, we generated images reflecting combinations of every pair of objects (drawing one from each list) under three different degrees of rotation (clockwise 90°, counterclockwise 90°, and 180°). This mirrors the stimulus generation approach used in Boger and Firestone (2025), though those stimuli were generated to vary real-world size with orientation (rather than animacy).

Several thousand images are generated as a result and then sorted to select anagrams that were uniquely identifiable as both prompts (before running any experiments). This yielded 12 images appearing in 6 anagram pairs: dog-boot, duck-airplane, elephant-sofa, sheep-car, rabbit-shoes, and horse-boat. Stimuli are available in our OSF repository. This approach controls for nearly all lower-level covariation associated with changes in animacy, allowing for more exacting control than is offered by conventional approaches. It also controls for relevant mid-level features that remain constant with orientation (e.g., configural shape, curvature, etc., which in some contexts are sufficient to produce differences in perceived animacy; e.g., Long et al., 2017; Schmidt et al., 2017; Zachariou et al., 2018; Yetter et al., 2021) — and our specific designs even preserve the relevant ensemble statistics of the displays across trial types. (The anagram pairs are pixelwise-identical subject to rotation, which means that orientation and aspect ratio can change when rotating the anagrams. However, Experiment 7 ensured that such changes cannot account for our observed effects.)

### Experiments 1–2: Visual working memory

Experiments 1–2 asked whether animacy affects visual working memory, using a change detection task. Our key question was whether changes that alter an object’s animacy (e.g., replacing a rabbit with a boot) are more detectable than changes that do not alter an object’s animacy (e.g., replacing a rabbit with a dog — even when the dog and the boot are the same image, just rotated). The pre-registration for Experiment 1 is available at https://aspredicted.org/uu7r5t.pdf; the pre-registration for Experiment 2 is available at https://aspredicted.org/km4nj2.pdf.

#### Stimuli and procedure

Participants were instructed to remember an array of objects and then detect any changes to the array. On each trial, the initial array contained 5 unique objects, such that (a) no anagram pairs repeated within an array and (b) one anagram pair could be held out as the ‘changing’ object. This initial array was visible for 1000ms, after which it disappeared and a 3000ms retention interval began. Following this retention interval, a new array appeared, where one object from the initial array changed identity. The array remained on screen until the participant detected which object changed. Participants responded by first pressing the space bar upon locating the object that changed, and then using the mouse to click on the location of that object.

Participants completed 120 such trials. Each of 6 anagram pairs (i.e., 12 objects) underwent a cross-category (i.e., animacy altered) change 5 times and a within-category (i.e., animacy unaltered) change 5 times. Trial order was randomized for each participant.

The task also contained 8 catch trials which depicted circles of varying colors (instead of the anagrams stimuli). 4 catch trials appeared at the start of the task, and 4 were randomly interspersed throughout.

Experiment 1 was identical to Experiment 2, with one key exception. In Experiment 1, before the task began, participants completed a ‘labeling’ phase, in which they viewed a gallery of the anagrams used in the task, were shown candidate labels for those images, and had to click on the images corresponding to those labels. This ensured that participants correctly identified each anagram stimulus as its intended object. Experiment 2 did not contain this labeling phase, which (a) allowed us to examine whether the anagrams are readily identifiable without explicit prompting, and (b) ensured that the labeling phase did not bias participants.

#### Analysis and results

We recruited 30 participants each for Experiments 1–2. In Experiment 1, 2 participants were excluded due to our pre-registered accuracy criterion (which excluded participants who failed to respond correctly on at least 30% of catch trials); and 194/3584 (5.4%) of trials were excluded for having response times which were either too slow (above 5000ms) or too fast (below 300ms).

We observed higher accuracy for cross-category changes than for within-category changes (*M* = 3.7%, *t*(27) = 2.36, *p* = 0.025, *d* = 0.45), suggesting that visual working memory encodes animacy itself.

In Experiment 2, 2 participants were excluded for accuracy, and 223/3584 (6.2%) of trials were excluded for speed. Here, we replicated the effect observed in Experiment 1: Participants were more accurate on cross-category than within-category changes (*M* = 4.7%, *t*(27) = 2.99, *p* < .01, *d* = 0.57).

### Experiments 3–4: Visual search

Experiments 3–4 examined whether animacy guides more basic visual processing. The pre-registration for Experiment 3 is available at https://aspredicted.org/i85iv9.pdf; the pre-registration for Experiment 4 is available at https://aspredicted.org/aq2tw5.pdf.

#### Stimuli and procedure

Participants completed a visual search task modeled after Long et al. (2017). On each trial, a target object was previewed for 1000ms. The target then disappeared and there was a 500ms inter-stimulus interval (ISI), after which 6 objects appeared, radially positioned around a central fixation cross. The 6 objects each depicted a unique anagram (i.e., none of the objects within a given trial were anagrams of each other). Participants had to search for the previewed target object (which was always among the 6 objects in the search array). Upon finding the target, participants pressed the space bar, after which point all the images were replaced with numbers (arranged 1 through 6, clockwise). Participants used their keyboard to select the number that occupied the location of the target.

Participants completed 144 test trials (in a random order for each participant). Each of 6 anagram pairs (i.e., 12 objects) appeared as the target in each of the 6 locations (1 through 6) twice, once in a “mixed animacy” trial and once in a “uniform animacy” trial. In mixed animacy trials, the target differed in animacy from its distractors (i.e., animate target surrounded by inanimate distractors, or inanimate target surrounded by animate distractors); in uniform animacy trials, the target did not differ in animacy from its distractors (i.e., animate target surrounded by animate distractors, or inanimate target surrounded by inanimate distractors). Additionally, 8 catch trials were included (depicting basic shapes) — 4 at the start of the experiment, and 4 randomly interspersed throughout.

Experiment 3 contained the labeling phase described in Experiment 1, while Experiment 4 did not; otherwise, the two experiments were identical.

#### Analyses and results

Participants were excluded for failing to respond correctly on at least 75% of catch trials; and trials with response times below 300ms or above 3000ms were excluded. In Experiment 3, 30 participants completed the task, 1 of which was excluded (and 222 of 4408, or 5.0%, of trials were excluded); in Experiment 4, 30 participants completed the task, none of which were excluded (and 165 of 4560, or 3.6%, of trials were excluded).

In both experiments, we observed faster response times for mixed animacy arrays than uniform animacy arrays (Experiment 3: *M* = 108ms, *t*(28) = 7.04, *p* < .001, *d* = 1.31; Experiment 4: *M* = 124ms, *t*(29) = 7.10, *p* < .001, *d* = 1.30). Additionally, this effect did not arise merely because of a speed-accuracy tradeoff, as participants were no less accurate on mixed animacy trials (and if anything were numerically more accurate; Experiment 3: *M* = 0.8%, *t*(28) = 1.34, *p* = 0.19, *d* = 0.25; Experiment 4: *M* = 0.7%, *t*(29) = 1.05, *p* = 0.30, *d* = 0.19).

### Experiments 5–6: Visual search (present/absent)

While Experiments 3–4 tested effects of animacy on visual search with a task used in previous work (Long et al., 2017), Experiments 5–6 asked whether such effects might arise in a more typical visual search task. Here, we presented participants with a present/absent task. The pre-registration for Experiment 5 is available at https://aspredicted.org/bn5qj9.pdf; the pre-registration for Experiment 6 is available at https://aspredicted.org/5k7ig2.pdf.

#### Stimuli and procedure

On each trial, a target object was previewed for 1000ms. Following a 500ms ISI, a search array appeared, presenting 5 objects (randomly positioned in an invisible 3×4 grid). Participants simply had to say whether the target object was present or absent in this array. The task contained 96 test trials (in a randomized order for each participant), equally split between target-present trials and target-absent trials. The 48 target-present trials were themselves equally split, with one half depicting mixed animacy arrays (where the target differs in animacy from its distractors), and the other half depicting uniform animacy arrays (where the target does not differ in animacy from its distractors). Thus, each of 6 anagram pairs (12 objects) was the previewed target in 4 target-absent trials, 2 target-present-mixed-animacy trials, and 2 target-present-uniform-animacy trials.

As before, the task began with 4 catch trials (and 4 additional catch trials were randomly placed throughout the task). Experiment 5 differed from Experiment 6 only in that the former contained the labeling phase described in Experiment 1, while the latter did not.

#### Analyses and results

Here, we used the same exclusion criteria as in Experiments 3–4: Participants were excluded for responding correctly on less than 75% of catch trials (which excluded 0 participants in Experiment 5 and 1 participant in Experiment 6), and trials with response times below 300ms or above 3000ms were excluded (excluding 38 of 3120 trials in Experiment 5 and 45 of 3016 in Experiment 6).

As in our previous search experiments, search was faster on mixed animacy arrays than uniform animacy arrays (Experiment 5: *M* = 94.4ms, *t*(29) = 5.37, *p* < .001, *d* = 0.98; Experiment 6: *M* = 78.5ms, *t*(28) = 4.80, *p* < .001, *d* = 0.89). Similarly, participants were significantly more accurate on mixed animacy arrays in Experiment 5 (*M* = 3.3%, *t*(29) = 2.69, *p* = .01, *d* = 0.49), and numerically so in Experiment 6 (*M* = 0.5%, *t*(28) = 0.36, *p* = 0.72, *d* = 0.07).

### Experiment 7: Controlling for orientation

Experiment 7 ensured that differences in orientation cannot explain our previously observed effects of animacy on visual attention. In other words, even though the animate and inanimate versions of a visual anagram contain the same pixels, they differ in orientation — and perhaps that difference accounts for our results. This concern is especially acute in the case of visual search, where even very minor visual differences allow targets to stand out from distractors. To address this, we replicated the present/absent search task from Experiments 5–6, but this time included trials that depicted silhouetted versions of the anagrams. If such trials do not show a search advantage (while the anagram trials do), then orientation alone cannot explain our results. The pre-registration for Experiment 7 is available at https://aspredicted.org/hk9e52.pdf.

#### Stimuli and procedure

We generated silhouetted versions of the visual anagrams by converting them into convex hulls with OpenCV. Thus, the silhouetted stimuli retain the aspect ratio and orientation of the original images, but they present as indistinct blobs that are not readily identifiable; in this way, they test for effects of orientation without animacy.

We used the same task timing and design as in Experiments 5–6. However, in this experiment, the 96 test trials were split up differently. One half of trials depicted visual anagrams (as in Experiments 5–6), and the other half depicted silhouetted versions of those anagrams (within-subjects). The two sets of trials were separated into blocks, and block order was randomized for each subject. Each block (of 48 trials each) was then equally split to contain 24 target-absent trials, 12 being target-present-mixed-animacy trials, and 12 target-present-uniform-animacy trials. This design allowed us to (a) replicate our effects from Experiments 5–6, (b) ask whether similar effects arise with silhouettes, and (c) directly compare the two effects in a statistically powerful within-subjects design.

#### Analyses and results

We recruited 150 participants for this experiment, 149 of which completed the task. Then, we applied the same exclusion criteria as in Experiments 5–6, which excluded 4 participants and 292 of 15080 trials (1.9%).

First, we found that the mixed-animacy search advantage for anagrams replicated again (*M* = 75.7ms, *t*(144) = 6.55, *p* < .001, *d* = 0.54). Second, we found *no* search advantage for the silhouetted stimuli (*M* = 2.16ms, *t*(144) = 0.17, *p* = 0.86, *d* = 0.01, BF10 = 0.09). Finally, and perhaps most importantly, the difference in search advantage between the two stimulus classes was significant (*M* = 73.5ms, *t*(144) = 4.24, *p* < .001, *d* = 0.35). Thus, changes in orientation or aspect ratio alone cannot explain our results.

As above, we examined differences in accuracy in these two conditions. Participants were no less accurate on mixed-animacy anagram trials (and if anything were numerically more accurate; *M* = 1.8%, *t*(144) = 1.88, *p* = 0.06, *d* = 0.16); and they were also numerically more accurate on mixed-animacy silhouette trials (*M* = 1.1%, *t*(144) = 0.98, *p* = 0.33, *d* = 0.08).

### Experiment 8: Controlling for semantic effects

Experiment 8 asked whether semantic processing alone (independently of perceived animacy) explains our previously observed results. Perhaps the anagram images merely activate a semantic label for animacy, thus explaining our results without necessarily appealing to visual processing of animacy itself. To address this, we used the exact same design as in Experiment 7, but replaced the silhouette trials with words trials (i.e., trials that presented linguistic labels for the anagrams). The pre-registration for Experiment 8 is available at https://aspredicted.org/73hj6t.pdf.

#### Stimuli and procedure

This design was identical to Experiment 7 in every respect, including trial numbers, block structure, etc. The only difference was that instead of containing one block of anagram trials and one block of silhouette trials, this experiment contained one block of anagram trials and one block of words trials. Words described the anagram images (e.g., the picture of a duck was labeled as “duck”) in an attempt to roughly match the word length of the two lists.

#### Analyses and results

We recruited 30 participants for this experiment. We applied the same exclusion criteria as in Experiment 7, which excluded 0 participants and 60 of 3120 trials (1.9%).

As in prior experiments, we observed a search advantage for anagrams (*M* = 114ms, *t*(29) = 5.04, *p* < 0.001, *d* = 0.92). However, we observed no search advantage for the words (*M* = −6.38ms, *t*(29) = 0.30, *p* = 0.77, *d* = 0.05, BF10 = 0.20); and the difference in search advantages between the anagrams and words was itself significant (*M* = 120ms, *t*(29) = 3.68, *p* < 0.001, *d* = 0.67). This suggests that semantic labeling cannot account for our results.

Participants were numerically more accurate on the mixed-animacy anagram trials (*M* = 0.6%, *t*(29) = 0.24, *p* = 0.82, *d* = 0.04). Participants were also numerically more accurate on the mixed-animacy words trials (*M* = 1.2%, *t*(29) = 0.51, *p* = 0.61, *d* = 0.09).

### Learning effects?

Upon the suggestion of a reviewer, we report exploratory analyses to ask whether learning or familiarity effects could explain our key findings. We analyze this in two ways. First, we ask whether our key effects arise early, by analyzing the first 10 trials of each type in each experiment (e.g., the first 10 mixed animacy vs. uniform animacy trials in the search experiments). Second, we ask whether the effects differ between the first 10 trials of each type and the last 10 trials of each type. We note that our experiments were not designed with the below analyses in mind, which may be underpowered for the questions they seek to answer. However, they do overall suggest that learning is not the explanation of our findings.

We found evidence that our key effects arise early. In 5/6 experiments (Experiments 1, 3, 4, 5, and 6), the effect was significantly (or in one case marginally) present even at the beginning of the experiment (Experiment 1, *p* = 0.07; Experiment 3, *p* = 0.01; Experiment 4, *p* < 0.001; Experiment 5, *p* < 0.001; Experiment 6, *p* < 0.01), and most of these results survive correction for multiple comparisons. In only one experiment (Experiment 2) was there a numerical disadvantage, but it was not significant (*p* = 0.34). (We refrain from analyzing Experiments 7–8 here because the concern of the learning effects does not apply to silhouettes or words, as those stimuli did not produce a search effect in the first place.)

Second, we found little to no difference between the start of the task and the end of the task. 4/6 experiments revealed numerically weaker effects at the end of the task than the start, though in each case the differences were not significant (Experiment 1, *p* = 0.63; Experiment 3, *p* = 0.76; Experiment 4, *p* = 0.57; Experiment 5, *p* = 0.08). 2/6 experiments revealed numerically stronger effects at the end of the task than the start, though only one was significant and would not survive a correction for multiple comparisons (Experiment 2, *p* = 0.01; Experiment 6, *p* = 0.23). Together, these analyses suggest that learning and familiarity are unlikely to drive our key effects.

